# iPSC-derived extracellular vesicles rescue deficits in human and mouse models of Parkinson’s disease

**DOI:** 10.64898/2026.05.19.726142

**Authors:** Wote Amelo Rike, Utkarsh Tripathi, Yara Hussien, Ashwani Choudhary, Aleksandar Rajkovic, Omveer Sharma, Idan Rosh, Andreea Manole, Fred H. Gage, Henry Houlden, Claude Brodski, Shani Stern

**Author notes:** Contributed equally (Wote Amelo Rike, Utkarsh Tripathi).

## Abstract

Parkinson’s disease (PD) pathogenesis often involves progressive α-synuclein (α-Syn)-mediated neuronal dysfunction, yet the earliest cellular events that link α-Syn pathology to circuit failure remain poorly defined. Here, we used human induced pluripotent stem cell (iPSC)-derived dopaminergic (DA) neurons from patients carrying the familial A53T SNCA mutation to reconstruct a temporal course of dysfunction *in vitro*. We identified a biphasic trajectory with an early phase of hyperexcitability, characterized by elevated spontaneous firing, followed by a progressive transition into hypoexcitability as the neurons mature, accompanied by reduced network activity, synaptic dysfunction, and α-Syn accumulation. Transcriptomic profiling at the critical transition point revealed a dual transcriptional signature, with upregulation of stress-inflammatory pathways (p53, JAK-STAT, apoptosis) coupled with systematic downregulation of metabolic and synaptic maintenance genes. This molecular profile preceded functional collapse, linking early hyperactivity-driven metabolic stress to subsequent neuronal exhaustion. To counteract this pathology, we used extracellular vesicles (EVs), small membrane-bound particles carrying intercellular signals, as a cell-free treatment approach. Strikingly, treatment with EVs derived from healthy iPSCs completely rescued both electrophysiological deficits and pathological α-Syn accumulation, restoring normal firing patterns, synaptic function, and network activity. Consistent with these observations, EV treatment reduced α-Syn aggregation and improved motor responses in α-Syn fibril-injected mice, which are characterized by pathological α-Syn accumulation and motor deficits. Overall, these findings demonstrate that EVs derived from healthy iPSCs can reverse PD-related phenotypes in human and mouse models.

## Introduction

Parkinson’s disease (PD) is the most prevalent synucleinopathy, followed by dementia with Lewy bodies (DLB) and multiple system atrophy (MSA) (1–4). It is also the fastest-growing neurological disorder in terms of mortality and disability (5), with its incidence and prevalence increasing with age, and the majority of diagnoses occurring after 60 (5, 6). As life expectancy rises and competing causes of death decline, the global PD burden is expected to grow, with cases projected to increase from 6.2 million in 2015 to 12.9 million by 2040 (7, 8). Pathologically, it is characterized by a loss of dopaminergic neurons (DA) in the substantia nigra and the presence of Lewy bodies (LBs) in the midbrain, leading to a progressive loss of motor control that manifests as bradykinesia, rigidity, resting tremor, and postural instability (9). However, the progression of PD varies widely, with nonmotor symptoms such as hyposmia, autonomic dysfunction, and REM sleep behavior disorder often emerging long before motor and cognitive impairments (10, 11). This progressive neurodegeneration, together with the delayed onset of clinical symptoms, makes early diagnosis difficult and complicates the development of effective therapy.

PD arises from a complex interplay between environmental and genetic factors, with most cases being idiopathic (12). In familial forms of the disease, several genetic mutations have been identified, including those in *SNCA*, *LRRK2*, *PARK2*, *PINK1*, *VPS35*, *GBA*, and *DJ-1* (*13, 14*) genes. α-Syn, encoded by the *SNCA* gene, has gained attention as a key player in the development of PD, with extensive clinical and experimental evidence supporting a causal link between environmental and genetic factors (15–17) α-Syn is a 14 kDa vertebrate-specific, highly soluble cytoplasmic protein primarily found in neurons, where it exists as an unstructured monomer in the cytosol in its native state (18, 19). Upon membrane binding, α-Syn undergoes a conformational change to form an α-helical structure and then multimerizes, promoting synaptic vesicle clustering and enhancing SNARE complex assembly essential for neurotransmission (20). Despite these functional roles, its precise physiological function is not fully understood. Under pathological conditions, α-Syn misfolds and aggregates into toxic oligomers and fibrils, which disrupt synaptic function, impair mitochondrial activity, and elicit neuroinflammatory responses, ultimately leading to neuronal loss (21). Notably, α-Syn can propagate between cells via EVs, contributing to the progressive degeneration of dopaminergic neurons and the spread of pathology in PD (22–24).

Although advances in cellular biology have opened new avenues for understanding PD, translating these discoveries into effective clinical treatments remains a complex and challenging journey. Current PD therapies mainly target motor symptoms using levodopa, frequently paired with dopamine agonists or MAO-B inhibitors to prolong benefits, but extended levodopa use may cause motor complications that worsen management over time (11). Deep Brain Stimulation (DBS) of the subthalamic nucleus can help in drug-resistant cases, yet neither approach addresses nonmotor symptoms or alters disease progression (25, 26). Consequently, growing research efforts are focusing on disease-modifying strategies that target various mechanisms, such as reducing α-Syn buildup, enhancing mitochondrial function, modulating neuroinflammation, and employing gene or cell-based therapies (27). Although several such approaches are now active with numerous Phase 1 and 2 trials, few disease-modifying therapies progress to Phase 3, and none have demonstrated clear evidence of slowing disease progression to be approved for clinical use (27–29). Given these limitations, there is a pressing need for novel therapeutic approaches.

Extracellular vesicles (EVs) have emerged as promising candidates due to their ability to modulate α-Syn pathology and support neuronal function in PD (22–24). These nanosized, spherical vesicles are enclosed by a single lipid bilayer and are secreted by nearly all cell types (30). Often described as replicas of their parent cells, EVs carry unique biomolecular signatures such as proteins, lipids, and nucleic acids that reflect the characteristics of their cells of origin (31). Stem cell-derived EVs (SC-EVs) exhibit therapeutic properties similar to those of their parent stem cells, including the promotion of anti-inflammatory responses, modulation of immune activity, and support for tissue repair (32–34). Thus, they can serve as a potent surrogate for stem cell therapy, even with better safety and feasibility, as stem cell therapy suffers from infusion toxicity, immunogenicity, tumorigenic potentials, and ethical issues (32, 35).

EVs derived from neural stem cells (NSC-EVs) have demonstrated significant neuroprotective effects in models of both neurodegenerative and neurodevelopmental disorders. In Alzheimer’s disease, NSC-EVs inhibit key pro-inflammatory signalling pathways, thereby reducing neuroinflammation and neuronal damage (30). A recent study using human iPSC-derived cortical neurons has also identified iPSC-derived EVs as modulators of neuronal function and potential therapeutic agents for autism spectrum disorder (ASD) (36). Furthermore, EVs from oligodendrocytes have been demonstrated to contribute to the maintenance of neuronal integrity by facilitating bidirectional neuron-glia communication and transferring antioxidant enzymes such as superoxide dismutase and catalase, which help mitigate oxidative stress (37, 38). In the context of PD, human NSC-derived EVs have shown promising therapeutic potential by attenuating oxidative stress, suppressing inflammation, and preventing dopaminergic neuronal loss in both *in vitro* and *in vivo* models (39). These findings highlight the potential of EVs to modulate key pathological processes in neurological disorders, supporting their development as a promising, disease-modifying therapeutic strategy for PD.

Although substantial evidence points to α-Syn pathology as a key driver and synaptic dysfunction as one of the earliest and most critical events in the pathogenesis of PD (40–42), effective therapeutic strategies targeting these mechanisms remain limited. Among emerging approaches, EVs derived from healthy cells have garnered interest for their ability to mediate intercellular communication and deliver functional bioactive cargo, including proteins, lipids, and RNAs, which can modulate neurodegenerative processes. Despite this promise, their ability to rescue functional deficits in patient-derived DA neurons and modify disease progression *in vivo* remains insufficiently explored. In this study, we employed human iPSC-derived DA neurons, carrying the PD-associated A53T mutation, to investigate neurophysiological alterations across developmental stages and to evaluate the therapeutic potential of EVs derived from healthy iPSC lines. We applied iPSC-derived EVs in both *in vitro* and *in vivo* models of PD and assessed their impact on disease-relevant phenotypes, including α-Syn accumulation, synaptic activity, and neuronal viability. This study seeks to promote the development of EV-based therapies as an innovative and disease-modifying approach for the treatment of PD.

## Methods

### Ethical consideration

All study participants provided informed consent, and ethical approval for the experiments in the study was obtained from the Institutional Review Board (IRB) of the University of Haifa. All the experiments were conducted per the Declaration of Helsinki and the regulations of the Israeli Ministry of Health.

### Characteristics of the cell lines

We derived three hiPSC lines from healthy individuals and three hiPSC lines from PD patients carrying the familial A53T mutation in the SNCA gene (Table 1). Each group contained two males and one female, and the average ages in each group were 54.3±1.5 (PD) and 55.3±8.3 (control).

**Table 1:** Clinical characteristics of the participants.

| Cell lines | Source | Sex | Age |
| --- | --- | --- | --- |
| UOHi033 | PD patient | M | 52 |
| UOHi034 | PD patient | F | 54 |
| UOHi035 | PD patient | M | 57 |
| UOHi036 | Control | M | 71 |
| UOHi008 | Control | F | 52 |
| UOHi037 | Control | M | 43 |

### Maintenance of iPSC Culture

Human iPSCs reprogrammed from patients and healthy controls were cultured in mTesR PLUS media. Cells were maintained in 6-well tissue culture plates and initially supplemented with the ROCK inhibitor Y-27632 for 24 h. The following day, cultures were fed with mTeSR PLUS medium without ROCK inhibitor. iPSCs were passaged when reaching 70-80% confluency using Gentle Cell dissociation reagent (100-0480, STEMCELL Technologies) for 6 minutes at room temperature. All iPSC lines were maintained at passages below 50 and were confirmed to have a normal karyotype.

### Dopaminergic (DA) differentiation

For the differentiation of iPSCs into DA neurons, we used the protocols described in Kriks et al (43) and also reported in our previous studies (40–42). Briefly, the iPSCs from patients and healthy controls were grown until ∼80% confluency. Then the cells were dissociated with Accutase (AT-104, STEMCELL Technologies) and re-plated at a density of 9×10^5^ cells per well (of a 6-well plate) on Matrigel-coated plates in mTesR medium (Stem Cell Technologies) with ROCK inhibitor Y-27632 (72308, STEMCELL Technologies). At 50% confluency (day 0 of differentiation), the media were replaced with knockout serum replacement (KSR) media containing DMEM F-12 (Biological Industries, 01-170-1A) with Glutamax (35050-038, Gibco), 15% knockout serum (KO-SR) (10828-010, Gibco), 1% non-essential amino acid (NEAA) (11140035, Gibco), 0.1LmM β-mercaptoethanol (219024280, MPB-MP Biomedicals), and supplemented with 100 nM LDN-193189 (1066208, Biogems) and 10LM SB431542 (Cayman Chemical Company, 0573051-151). For days 1–4, KSR medium containing 100 nM LDN193189, 10 μM SB431542, 0.25LM SAG (biogems, 9128694), 2 μM purmorphamine (biogems, 4831086), and 100 ng/mL FGF8b (100-25, Peprotech) was added daily, supplemented with 3µM CHIR99021 (biogems, 2520691) on days 3 and 4. From day 5 to day 10, KSR medium was gradually changed to N2 medium, composed of DMEM F-12 with Glutamax, 1% N2 supplement (17502-048, Gibco). For days 5 and 6, a mixture of 75% KSR (supplemented with all molecules in days 3 and 4) and 25% N2 medium was added to the cells, followed by equal amounts of KSR (supplemented with 100 nM LDN193189 and 3µM CHIR99021) and N2 media on days 7–8. On days 9–10, 25% KSRL(supplemented with small molecules in days 7-8)Lmedium was mixed with 75% N2 media, and then changed to B27 medium (Neurobasal medium (NB; 21103049, Gibco), 2% B27 supplement, 1% glutamax, 10 ng/mL BDNF, 10 ng/mL GDNF (PHC7044, Gibco), 1 ng/mL TGFβ3 (100-36E, Peprotech), 0.2 mM ascorbic acid (5088177, Biogems), and 0.1 mM cAMP (1698950, Biogems)) on day 11. The differentiation medium was replaced with a new medium each day until the second dissociation.

Between days 20 and 25 of the differentiation, neurons were dissociated and replated onto coverslips coated with Matrigel. They were then maintained in B27-containing medium to support further maturation until day 30. Starting from day 30, the B27 medium was progressively replaced with BrainPhys medium (Stem Cell Technologies, 05790) to facilitate the formation of synaptic connections.

### EVs extraction and treatment *in vitro*

EVs were isolated from iPSCs derived from a healthy donor using differential centrifugation followed by ultracentrifugation (44). Characterization was performed using a Zetasizer (Malvern Panalytical) to assess surface charge, particle size, and concentration via dynamic light scattering (DLS) and electrophoretic light scattering (ELS) techniques (36). The EV treatment was applied once a week starting from the tenth day of differentiation.

### Immunocytochemistry

Coverslips with DA neurons were fixed with 4% paraformaldehyde (PFA) for 15Lminutes at room temperature, followed by 3 washes with Dulbecco’s Phosphate-Buffered Saline (DPBS) (14287-072, Gibco). The cells were then permeabilized and blocked for one hour in DPBS containing 0.2% Triton X-100 and 10% Horse serum. Primary antibodies, α-Syn antibody (pSer129) (Biosciences, SMC-600D; 1:1000) and anti-MAP2 (Abcam, ab92434; 1:500) were diluted in blocking solution (DPBScontaining 0.2% Triton X-100 and 10% Horse serum) and applied to the cells. The cells were then incubated at 4° C overnight. The next day, the coverslips were washed three times with DPBS (5Lminutes each), incubated with the Alexa Fluor^TM^-conjugated secondary antibodies, and counterstained with DAPI (ab228549, Abcam) staining solution (1:2500) for 60Lminutes at room temperature. After final washes, the coverslips were mounted onto slides using Fluoromount-G mounting medium (Southern Biotech, 0100-01) and left to dry overnight in the dark. The fluorescence imaging was performed using a Nikon A1-R confocal microscope. The images were processed with NIS Elements 5.21 (Nikon) and analyzed using Imaris software version 9.8 (Oxford Instruments, Zurich, Switzerland).

### RNA extraction, sequencing, and analyses

At the seventh week post-differentiation, total RNA was extracted from 3 to 5 million DA neurons per sample using the Zymo RNA Clean & Concentrator kit (R1014) according to the manufacturer’s instructions. For every cell line, two biological replicates were grown separately. The high-capacity cDNA synthesis kit from AB Biosystems was used to reverse transcribe the isolated RNA.

RNA sequencing was performed on a NextSeq 500 sequencer, and the resulting reads were processed as FASTQ-format files for downstream analysis. The reads were trimmed with Trimmomatic, quality-checked with FASTQC v0.11.8, and aligned to the hg38 human genome with HISAT2. Gene-level counts were generated with featureCounts against the corresponding hg38 GTF gene annotation, producing a raw counts matrix across all samples. Differential expression (DE) analysis was conducted using the limma-voom(45) within Galaxy (Galaxy Training Network, “RNA-seq counts to genes (limma-voom)”). Briefly, low-abundance genes were filtered, library sizes were normalized using the Trimmed Mean of M-values (TMM), and voom precision weights were estimated before linear-model fitting with genotype as a factor. P values were adjusted by Benjamini–Hochberg correction to control for the false-discovery rate (FDR) (46). Unless noted otherwise, differentially expressed genes (DEGs) were defined as those with a false discovery rate (FDR) < 0.05 and an absolute logL fold change (**|**log_2_ fold change |) > 0.58 (≈1.5-fold). Enrichment analyses for KEGG and GO terms were performed in SRPlot based on the significant DEGs.

### Electrophysiological Recordings

Whole-cell patch-clamp recordings were conducted on DA neurons at weeks 5, 7, and 9 post-differentiation. For each cell line, 2-5 coverslips were recorded from several differentiation rounds. The same recording protocol was subsequently applied following differentiation with EV treatment. Recordings were performed at room temperature using a recording chamber filled with HEPES-based artificial cerebrospinal fluid (ACSF) containing (in mM): 10 HEPES (Sigma-Aldrich, Cat #H0887-100ML), 4 KCl (Sigma-Aldrich, Cat #P3911), 139 NaCl (Sigma-Aldrich, Cat #S9888), 2 CaCl2 (Sigma-Aldrich, Cat #223506), 1MgCl2 (Merck, Cat #7791-18-6) and 10 D-glucose (Sigma-Aldrich, Cat # G7021). The pH of the ACSF was adjusted to 7.4, and the osmolarity was adjusted to 310 mOsm. The recording micropipettes with a tip resistance of 10–15LMΩ were filled with an internal solution containing (in mM) K-gluconate (130LmM), KCl (6LmM), NaCl (4LmM), Na-HEPES (10LmM), K-EGTA (0.2LmM), GTP (0.3LmM), Mg-ATP (2LmM), cAMP (0.2LmM), d-glucose (10LmM), biocytin (0.15%), and rhodamine (0.06%) adjusted to a pH level of 7.4 and osmolarity to 290–300LmOsm. Data were acquired at a sampling rate of 20 kHz. Recordings were inspected for stability and signal quality by the series resistance and holding currents.

### Electrophysiological Data Analysis

#### Total evoked action potentials

The cells were recorded in current-clamp mode while maintaining a steady membrane potential of −60LmV using a constant holding current at three developmental time points. Neurons were subjected to depolarizing current injections consisting of 38 incremental steps (each step increasing by 3 pA, 400 ms duration per step). We quantified the total number of evoked action potentials (APs) across all steps (40).

#### Action potential shape analysis

We also performed detailed spike-shape analysis on the first evoked AP from each neuron that was generated with the least amount of injected current. The amplitude of the 5Lms fast after-hyperpolarization (fAHP) was calculated by measuring the difference between the spike threshold and the membrane potential recorded 5Lms after the voltage trace repassed the threshold level during the repolarization phase of the action potential. The spike amplitude was calculated as the difference between the maximum membrane potential reached during a spike and its threshold. The action potential width (full width at half maximum, FWHM) was defined as the duration between the ascending and descending phases of the spike at 50% of its amplitude. The spike threshold is defined as the membrane potential at which depolarization rapidly accelerates, marked by the first peak in the second derivative of the voltage-versus-time curve, signaling the initiation of an action potential (40–42).

#### Spontaneous action potential analysis

Spontaneous firing activity was quantified using custom Python scripts that automatically detected action potentials (spikes). To ensure accurate detection and exclude noise, spikes were identified using two criteria: a minimum peak height above -50 mV and a minimum prominence of 20 mV, meaning the spike had to stand out clearly from the surrounding signal. Bursts were defined as groups of at least three spikes occurring in rapid succession, with inter-spike intervals less than 0.5 s. From these recordings, multiple parameters were extracted, including total spike count, burst rate (bursts/minute), average burst duration (average time from first spike to the last spike within a burst), total burst time (sum of durations of all bursts in the recording), and average spike amplitude, enabling quantification of both overall neuronal activity and firing pattern dynamics.

#### Analysis of synaptic activity

We recorded spontaneous excitatory postsynaptic currents (sEPSCs) in control and A53T DA neurons at three developmental time points at a holding potential of –70 mV. sEPSCs were detected using a custom MATLAB algorithm (EPSCs_remove_Noise6.m) that processed raw traces through low-pass filtering (0.1 Hz), detrending, and 50 Hz noise removal. Events were identified as negative-going peaks exceeding 2.5× robust standard deviation, validated by exponential decay fitting (R² ≥0.7), and analyzed for amplitude, frequency, rise time, decay time constant (τ), and charge transfer. Event rates were calculated by dividing total detected events by recording duration, with inactive cells assigned zero values.

#### Maturation Index

A comprehensive maturation index was calculated for each group (control, A53T, and EV treatment) by normalizing the electrophysiological parameters across all conditions and timepoints using min-max scaling. Parameters positively associated with neuronal maturation (spike count, AP amplitudes, sEPSC amplitude, and sEPSC frequency) were directly normalized, such that higher values corresponded to greater maturity. In contrast, parameters where lower values indicate greater maturation (AP half-width and rise time) were normalized such that lower values yielded higher maturation scores. For spike threshold, normalization was directionally adjusted so that more negative values received higher maturation scores. The resulting index ranged from 0 (least mature) to 1 (most mature).

#### Z-Score Analysis

For comparative visualization, z-score normalization was applied to all parameters using z = (x -μ) / σ, where x is the individual value, μ is the population mean, and σ is the population standard deviation across conditions and timepoints. Features were ordered by A53T z-score values for optimal visualization, with below-average performance (negative z-scores) shown in light shades and above-average performance (positive z-scores) in dark shades.

### Animals

Male WT C57BL/6JRccHsd mice, aged 8-12 weeks, were purchased from Envigo, Israel. All animals were maintained in a temperature-controlled environment (21–23°C) under a 12-hour light/dark cycle, with unrestricted access to food and water in a pathogen-free facility. All procedures and experimental protocols involving the animals received approval from the Institutional Animal Care and Ethics Committee at Ben-Gurion University of the Negev (Permit number: IL-22-04-21-C).

### Stereotaxic surgery

Five-month-old male C57BL/6 RccHsd mice were deeply anesthetized using a combination of ketamine and xylazine and subsequently placed in a stereotactic headframe (Stoelting, IL, USA). Stereotactic injections were performed as previously described (47). A midline incision was made in the scalp, followed by the drilling of a burr hole in the appropriate location for targeting the striatum on the right side of the skull. α-Syn PFFs or monomers were injected at a total dose of 5 μg in 2.5 μL per animal, delivered at a rate of 0.25 μl/minute, using a 30-gauge needle connected to a 25 μL syringe (702 RN, Hamilton Company). The needle was left in place for an additional five minutes before being carefully withdrawn from the brain. According to the mouse brain atlas by Paxinos and Franklin, the injections were precisely targeted in the striatum at the following coordinates: anterior-posterior (AP) +0.4 mm, medial-lateral (ML) -2.0 mm relative to bregma, and dorsal-ventral (DV) -2.9 mm from the dural surface.

### EVs treatment -*in vivo*

A total of 12 adult mice were randomly assigned to one of three experimental groups, with four mice per group. The first group, serving as the control, received α-Syn monomers, while the second and third groups were injected with α-Syn PFFs, all into the striatum. The second group served as an untreated α-Syn PFFs control, whereas the third group received intranasal EV treatment starting after the α-Syn PFFs injection. EVs were administered every other day for 3.5 months, with each mouse receiving 4 µL containing 2–3 × 10¹L particles/µL per dose.

### Behavioral Test

#### Inverted screen test

The mice were initially placed in the center of a grid screen. After 2 seconds, the screen was inverted, causing the mice to hang upside down above a padded surface. The duration for which each mouse remained suspended was recorded, with a maximum observation period of 120 seconds. Three measurements were taken for each mouse, and the average was used for statistical analysis (48).

### Immunohistochemistry

Coronal brain sections containing the substantia nigra were cut at a thickness of 14 µm using a cryostat and stored at −80 °C until further processing. Dual immunofluorescence staining was performed on the same sections to visualize tyrosine hydroxylase (TH) and phosphorylated α-Syn at serine-129 (pS129-α-syn), for direct comparison of dopaminergic neurons and pathological α-Syn inclusions within the same tissue.

For immunostaining, sections were thawed at room temperature, fixed in cold acetone (−20 °C, 10 minutes), rinsed in PBS, and subjected to antigen retrieval in citrate buffer (pH 6.0) in a 90 °C water bath, followed by three washes in PBS (5 minutes each). Sections were then blocked for 1 hour at room temperature using normal goat serum (NGS) and bovine serum albumin (BSA). The blocking solution was discarded without washing, and sections were immediately incubated overnight at 4 °C with primary antibodies: rabbit anti-TH (AB152, Millipore; 1:500) and mouse anti-pS129-a-Syn (clone pSyn#64, FUJIFILM Wako Pure Chemical Corporation, 015-25191; 1:1000).

The following day, sections were washed three times in PBS and incubated for 1.5 hours at room temperature with secondary antibodies diluted 1:500 in PBS: Alexa Fluor® 647-conjugated goat anti-rabbit IgG (Jackson ImmunoResearch, 111-605-003) and Alexa Fluor® 488-conjugated goat anti-mouse IgG (Jackson ImmunoResearch, 115-545-003). After additional PBS washes, sections were mounted using Immumount medium containing DAPI for nuclear counterstaining and allowed to dry for 24 hours at room temperature before imaging.

### Statistical analysis

Electrophysiological recordings were acquired using Clampex v11.1. Raw data were processed using Clampfit 10 and analyzed with custom-written scripts in MATLAB (release 2014b; The MathWorks, Natick, MA, USA), following previously described methods (49). Graphs and statistical analyses were performed using GraphPad Prism (version). For the comparison of evoked action potentials, EPSC rates, and amplitudes, the Mann–Whitney *U* test was employed. All data values were presented as meanL±Lstandard error (SE). A value of *p*L<L0.05 was considered significant for all statistical tests. For RNA sequencing analysis, the FDR-adjusted mean value was evaluated, and FDRL<L0.05 was considered significant.

## Results

### Biphasic changes in neural excitability with sustained impairment of synaptic connectivity in A53T DA neurons compared to controls

To characterize how the A53T mutation influences neuronal maturation and intrinsic electrophysiological properties throughout the differentiation process (Fig. 1a), we performed whole-cell current-clamp recordings at three defined timepoints: week 5 (early differentiation) (Fig. 1b-d), week 7 (mid-stage differentiation) (Fig. e-g), and weeks 9–10 (late-stage differentiation) (Fig. 1h-j) post-differentiation. Then, we quantified the excitability by measuring the total number of evoked action potentials (APs) in each timepoint over all depolarization steps (see Methods). At the earliest differentiation timepoint (week 5), A53T mutant neurons were hyperexcitable compared to control neurons. Representative traces of evoked action potentials for control and mutant neurons are presented in Fig. 1b and c, respectively. The average number of evoked action potentials was significantly larger in A53T mutant neurons compared to control neurons (Fig. 1d) (P = 0.0006). By week 7, as shown with representative traces of evoked action potentials (Fig.1e, f), overall spike output in A53T mutant neurons was normalized, becoming comparable to controls (Fig. 1g). Notably, control neurons demonstrated a substantial maturation-driven increase in firing (a 142% rise compared to week 5), whereas A53T mutant neurons showed only modest enhancement (approximately 41%). The final timepoint (weeks 9–10) examined revealed the full manifestation of A53T-mediated pathology, with control neurons achieving mature, stable excitability profiles (Fig. 1h), while A53T mutant neurons underwent marked excitation reduction (Fig. 1i). This divergence was immediately apparent in firing capacity, with controls generating significantly more evoked action potentials than A53T mutant neurons (P = 0.002; Fig. 1j). These changes represent not merely delayed maturation but active regression toward a less mature, dysfunctional state. In contrast, control neurons maintained their mature profile with stable and developing electrophysiological parameters.

**Figure 1:**
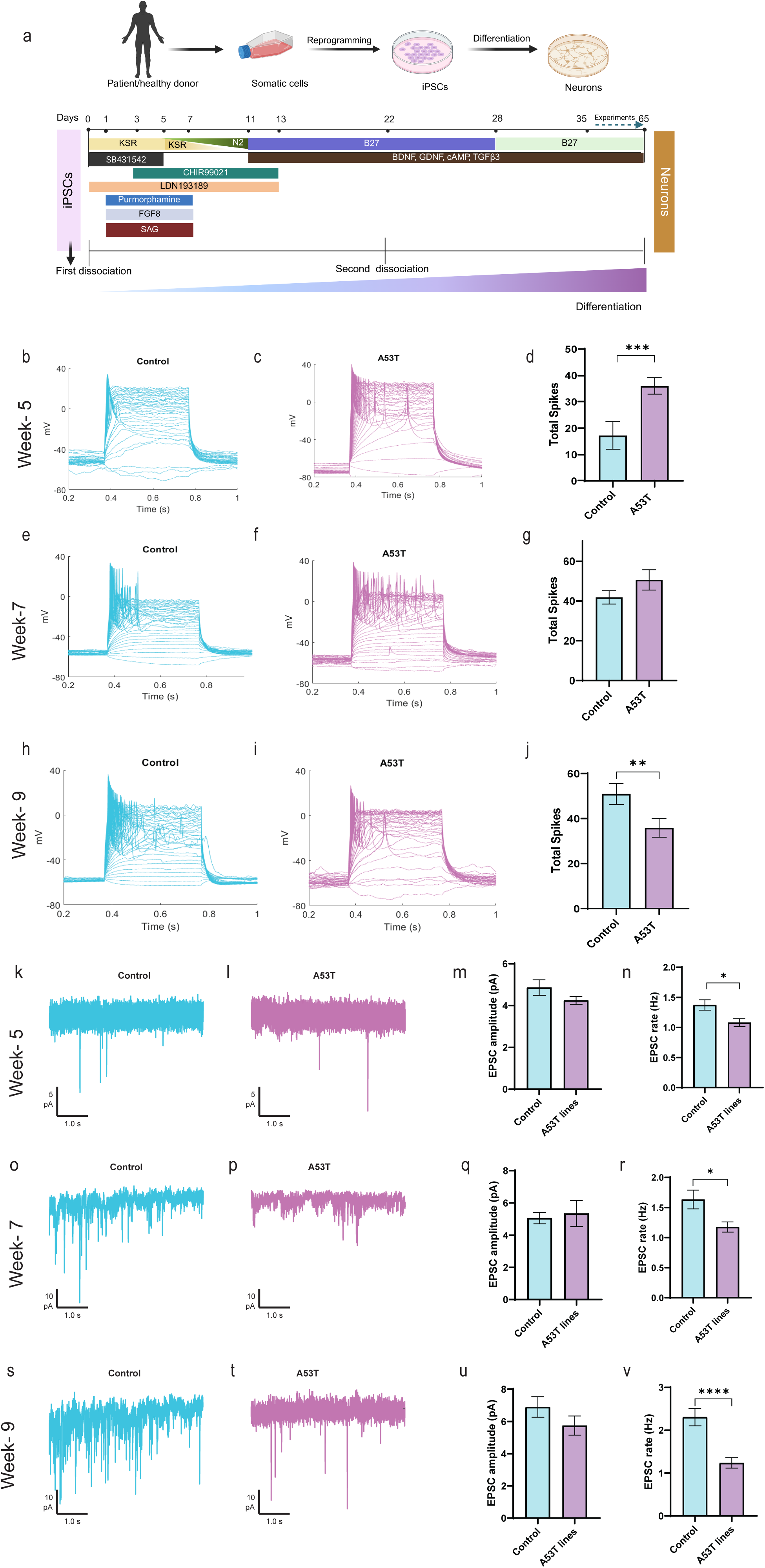
Biphasic Intrinsic Excitability with Persistent Synaptic Connectivity Deficits in A53T Dopaminergic Neurons Across Maturation. The familial A53T SNCA mutation imposes a biphasic program of intrinsic excitability in human DA neurons. Representative current-clamp recordings and quantitative analyses of evoked action potentials in control and A53T dopaminergic neurons at three developmental timepoints. **(a)** Schematics showing the differentiation of iPSCs into DA neurons. **Week 5** -Early maturation: Representative voltage traces showing evoked action potentials in **(b)** control (cyan) and **(c)** A53T mutant neurons (purple). **(d)** Quantification of total evoked spikes (Control: n=20; A53T: n=54). **Week 7** -Mid-stage maturation: Representative voltage traces showing evoked action potentials in **(e)** control (cyan) and **(f)** A53T mutant neurons (purple). **(g)** Quantification of total evoked spikes showing convergence of spike output. **Week 9**: Late maturation: Representative voltage traces showing evoked action potentials in **(h)** control (cyan) and **(i)** A53T mutant neurons (purple). **(j)** Quantification of total evoked spikes showing progressive decline in A53T excitability with reduced spike output (Control: n=72; A53T: n=69). Whole-cell voltage-clamp recordings of spontaneous excitatory postsynaptic currents (sEPSCs) at a holding potential of -60 mV. **Week 5**: Representative sEPSC traces from **(k)** control and **(l)** A53T mutant neurons. Quantification of sEPSC **(m)** amplitude and **(n)** frequency displaying a significant reduction in A53T mutant neurons compared to control (Control: n=17; A53T: n=57). **Week 7:** Representative sEPSC traces from **(o)** control and **(p)** A53T mutant neurons. Quantification of sEPSC **(q)** amplitude and **(r)** frequency showing persistent frequency deficits (Control: n=96; A53T: n=90). **Week 9**: Representative sEPSC traces from **(s)** control and **(t)** A53T mutant neurons. Quantification of sEPSC **(u)** amplitude and **(v)** frequency showing exacerbated synaptic dysfunction with severely reduced frequency (Control: n=74; A53T: n=65). Data represent mean ± SEM with statistical analysis by the Mann-Whitney U test. (*P<0.05, **P<0.01, ***P<0.001).

To investigate the synaptic consequences of the A53T SNCA mutation, we recorded spontaneous excitatory postsynaptic currents (sEPSCs) in control and A53T DA neurons at three developmental stages. Even at the earliest developmental stage (week 5) (Fig. 1k, l), A53T mutant neurons displayed synaptic impairments that would presage more severe dysfunction at later timepoints. While basic synaptic strength appeared intact, as evidenced by comparable sEPSC amplitudes between genotypes (4.9 ± 0.4 pA in control neurons vs. 4.3 ± 0.2 pA in A53T mutant neurons; n.s.; Fig. 1m), A53T mutant neurons nevertheless exhibited a 20% reduction in sEPSC frequency compared to controls (1.1 ± 0.07 Hz in A53T mutant neurons vs. 1.4 ± 0.09 Hz in control neurons; P = 0.015; Fig. 1n). These findings suggest impaired establishment or maintenance of excitatory synaptic connections during this critical developmental window. The intermediate developmental stage (week 7) revealed a persistent dysfunction in A53T mutant neurons (Fig. 1o, p). Synaptic strength continued to show no significant impairment, with sEPSC amplitudes remaining comparable between genotypes (5.1 ± 0.4 pA in control neurons vs. 5.3 ± 0.8 pA in A53T mutant neurons; n.s.; Fig. 1q). However, the sEPSC frequency decrease first observed at week 5 persisted, with A53T mutant neurons maintaining reduced sEPSC frequency (1.2 ± 0.1 Hz in A53T mutant neurons vs. 1.6 ± 0.2 Hz in control neurons; P = 0.03; Fig. 1r), confirming ongoing impairments in synaptic network formation. Importantly, this deficit progressively worsened at advanced stages of neuronal maturation (week 9-10), and the divergence in sEPSC frequency between control and mutant neurons became more pronounced (Fig. 1s, t). While sEPSC amplitude remained statistically comparable further decrease in sEPSC frequency was observed between the genotypes (5.7 ± 0.6 pA in A53T mutant neurons vs. 6.9 ± 0.6 pA in control neurons; n.s.; Fig. 1u). The frequency deficits that characterized A53T mutant neurons throughout development was more severe at week 9–10, with mutant neurons exhibiting less than half the sEPSC frequency compared to controls (1.2 ± 0.1 Hz in A53T mutant neurons vs. 2.3 ± 0.2 Hz in control neurons; P < 0.0001; Fig. 1v). This dramatic reduction indicated failure of synaptic network maintenance or progressive loss of functional excitatory inputs.

### Progressive suppression of spontaneous firing and burst activity in A53T DA neurons compared to healthy controls

To determine whether the intrinsic and synaptic excitability deficits described above (Fig. 1) translate into altered network-level dysfunction, we characterized spontaneous activity patterns by recording spontaneous action potentials at a fixed voltage clamp membrane potential (–45 mV) over 50 s recordings. This analysis encompassed spontaneous spike frequency, burst architecture (a burst is defined as more than 3 spikes with inter-spike intervals less than 0.5 s), burst duration, and spike amplitude characteristics across the developmental timeline. At early stage (week 5), while healthy control neurons were essentially silent with no spontaneous firing (0 spikes; Fig. 2a shows an example raster plot), mutant neurons exhibited robust spontaneous firing (0.16 ± 0.05 Hz; P < 0.0001 vs. controls; Fig. 2d). Bursts were detected exclusively in mutant neurons, with individual burst events lasting 1.0 ± 0.2 s; however, the overall time spent in bursting was low (0.13 ± 0.06 s) (Fig. 2e, f), indicating that the bursting occurred infrequently. Mutant neurons also displayed spontaneous firing with amplitude 55 ± 3 mV (Fig. 2g), and an average burst rate of 0.0028 ± 0.0012 Hz (Fig. 2h). These observations collectively identify a premature emergence of network-level hyperactivity specifically associated with the A53T mutation during initial neuronal network assembly.

**Figure 2:**
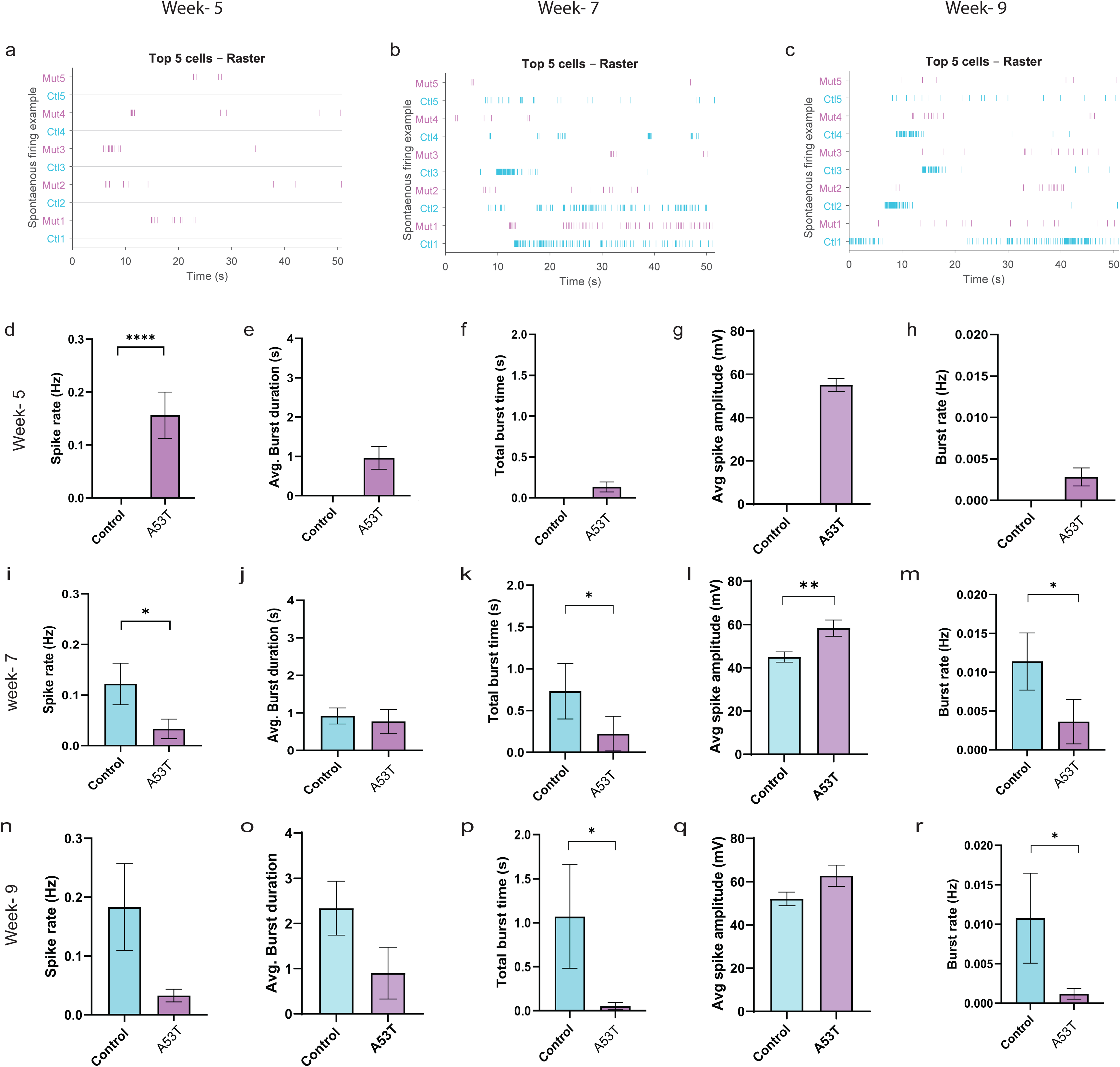
Altered Spontaneous Network Activity in A53T mutant neurons. Spontaneous firing and burst activity in A53T DA neurons emerge early, followed by impaired maturation. Representative raster plots showing spontaneous action potentials in the five most active neurons per condition in three timepoints. **Week 5: (a)** A53T mutant neurons exhibit premature hyperactivity while controls remain silent. **Week 7: (b)** Control neurons develop normal spontaneous activity, while A53T mutant neurons show declining activity. **Week 9: (c)** Markedly impaired spontaneous firing in A53T mutant neurons. Other measured electrophysiological parameters at the respective time points are as follows. **Week 5:** Compared to control, A53T mutant neurons exhibited **(d)** significantly increased spike rate, accompanied by a non-significant change in **(e)** average burst duration, **(f)** total burst time, **(g)** average spike amplitude, and **(h)** burst rate. **Week 7**: Compared to control, A53T showed **(i)** reduced spike rate, **(j)** unchanged average burst duration. **(k)** reduced total burst time. **(l)** increased average spike amplitude and **(m)** reduced burst rate. **Week 9**: Compared to control, A53T mutant neurons showed a non-significant change in **(n)** spike rate, **(o)** average burst duration, but showed a significant difference in **(p)** total burst time; **(q)** average spike amplitude remained unchanged, and **(r)** burst rate was significantly reduced. Sample sizes match those in Figure 1. Data represent mean ± SEM with statistical analysis by the Mann-Whitney U test. (*P<0.05, **P<0.01, ***P<0.001).

At the intermediate stage of synaptic maturation, control neurons developed robust spontaneous activity with significantly increased frequency of spikes (from 0 at week 5 to 0.128 ± 0.042 Hz) at week 7; Fig. 2b shows an example of raster plot), whereas A53T mutant neurons exhibited a substantial reduction in spontaneous firing from 0.16 ± 0.05 Hz at week 5 to 0.034 ± 0.020 Hz at week 7 (Fig. 2i). Although this difference between genotypes did not reach significance (P=0.08), it indicates a regressive trajectory in mutant neurons. Further analysis revealed significant genotype differences in burst-related parameters at week 7 where control neurons exhibited more frequent spontaneous bursting (P=0.034) than A53T mutant neurons (Fig. 2m). Moreover, the total burst time was markedly lower in mutant neurons (0.23 ± 0.21 s) compared to the control (0.73 ± 0.33 s) (P=0.036, Fig. 2k). Burst duration did not differ significantly between genotypes (0.91 ± 0.21 s, control vs. 0.77 ± 0.33 s in A53T mutant neurons; n.s., Fig. 2j); however, spike amplitudes remained significantly elevated in A53T mutant neurons relative to controls (58 ± 3.8 mV in A53T mutant neurons vs. 45 ± 2.4 mV in control neurons; P=0.004, Fig. 2l). These findings demonstrate that despite increased intrinsic spike amplitude, A53T mutant neurons fail to maintain early-established network activity, reflected by significant reductions in burst rate and total burst activity compared to controls at this intermediate stage.

At the latest stages (weeks 9–10), control neurons maintained robust spontaneous activity, slightly increasing their spike rate to 0.190 ± 0.076 Hz relative to week 7 (Fig.2c shows an example of raster plot), while A53T mutant neurons exhibited persistently low spontaneous firing (0.034 ± 0.012 Hz; P=0.11), showing no meaningful recovery and remaining well below control levels (Fig. 2n; see Fig. 2c for a representative raster plot). Analysis of burst dynamics at this advanced stage revealed substantial genotype differences. Control neurons displayed longer average burst durations (2.34 ± 0.60 s vs. 0.99 ± 0.57 s in A53T mutant neurons; Fig. 2o), though not statistically significant due to high variability. Importantly, total burst time was significantly reduced in A53T mutant neurons (0.05 ± 0.04 s in A53T mutant neurons vs. 1.07 ± 0.59 s in controls; P=0.044, Fig. 2p). Furthermore, the burst rate in A53T mutant neurons was significantly diminished compared to controls (0.0012 ± 0.0007 Hz vs. 0.0108 ± 0.0057 Hz) in A53T mutant neurons; P=0.049; Fig. 2r). Spike amplitudes no longer differed significantly (control: 52 ± 3 mV, A53T: 62 ± 5 mV; n.s., Fig. 2q), suggesting amplitude normalization at advanced stages.

Notably, comparing within-genotype patterns across timepoints highlighted distinct maturation trajectories: while control neurons progressively developed and maintained spontaneous firing, burst rate, and burst duration from week 5 to weeks 9–10, A53T mutant neurons exhibited an early abnormal peak at week 5 followed by a sustained regression in spontaneous activity, burst frequency, and total burst duration.

### A53T DA neurons exhibit concurrent cellular stress and inflammation-related signaling with metabolic suppression

To uncover the molecular mechanisms underlying the biphasic excitation and synaptic characteristics (Fig. 1, 2) observed in A53T mutant neurons, we performed RNA sequencing in control and A53T DA neuronal cultures at the critical week 7 timepoint, when functional deficits begin to emerge. Differential expression analysis identified a significant transcriptional shift between the control and A53T mutant neurons (|logL fold change| > 0.58, FDR < 0.05). A total of 1,190 differentially expressed genes (DEGs) were identified, of which 578 were upregulated and 612 downregulated (Fig. 3a).

**Figure 3:**
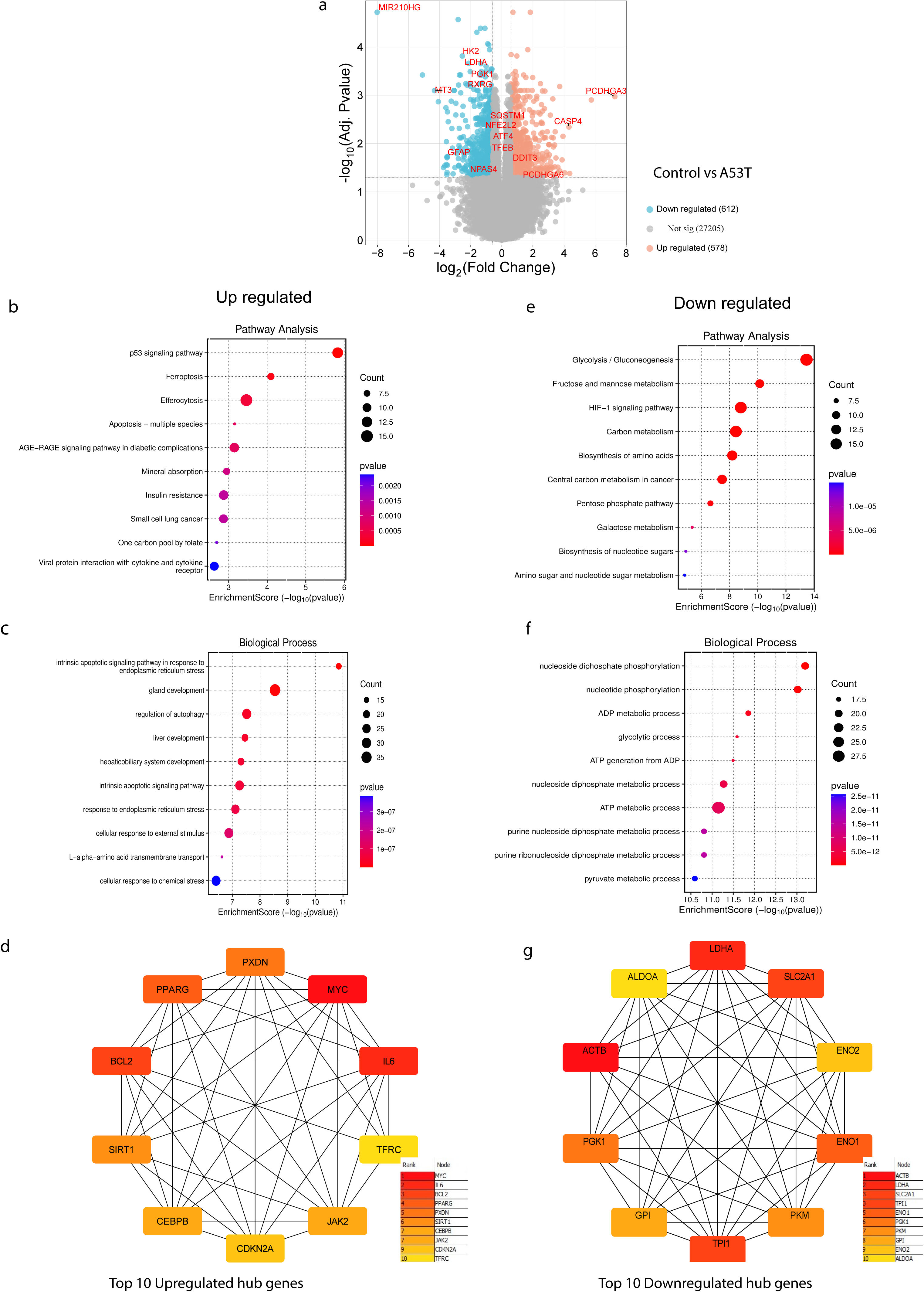
Transcriptomic Analysis Reveals Stress-Inflammation and Metabolic Dysfunction in A53T mutant neurons. RNA-seq reveals a dual “stress-inflammation” and “metabolic-shutdown” signature in A53T DA neurons. Differential expression analysis was performed at week 7 of differentiation**. (a)** Volcano plot showing global transcriptional changes with 578 upregulated (red) and 612 downregulated (blue) genes. Key genes labeled include stress markers (MIR210HG, ATF4, DDIT3) and metabolic enzymes (HK2, PGK1, LDHA). Upregulated pathways**: (b)** KEGG pathway analysis, **(c)** Biological Processes, **(d)** Network analysis of the top 10 hub upregulated genes showing central roles of IL6, PPARG, and other stress response regulators. Downregulated pathways **(e)** KEGG pathway analysis **(f)** Biological Processes **(g)** Network analysis of downregulated genes revealing metabolic enzyme hubs including ENO1, ACTB, and PGK1. Analysis based on n=3 control and n=3 A53T cell lines with 2 biological replicates each. DEGs were defined as those with an FDR < 0.05 and (**|**log_2_ fold change |) > 0.58 (≈1.5-fold).

The most enriched pathways among upregulated genes (Fig. 3b) were “p53 signaling pathway”, “ferroptosis”, “apoptosis”, and “AGE–RAGE signaling pathway in diabetic complications”, indicating activation of cellular stress-response and programmed cell death pathways. Gene Ontology (GO) biological process analysis (Fig. 3c) reinforced these findings, highlighting enrichment in “intrinsic apoptotic signaling pathway in response to endoplasmic reticulum stress”, “regulation of autophagy”, “responses to oxidative”, and “response to chemical stress”, consistent with activation of compensatory yet ultimately deleterious pathways. At the molecular function term level, enrichment of “NADH dehydrogenase activity” and “oxidoreductase-driven active transmembrane transporter activity” (Supp Fig. 1a) suggests a compensatory attempt to maintain redox balance. Cellular component analysis (Supp Fig. 1b) further showed membrane-associated alterations in A53T mutant neurons with enriched “basolateral plasma membrane “, “basal plasma membrane”, “organelle outer membrane”, and “collagen-containing extracellular matrix”. Network analysis identified central dysregulated hub genes spanning inflammatory (IL6), cell-cycle (CDKN2A, MYC), and survival-related (BCL2) pathways (Fig. 3d), underscoring coordinated stress-response signaling in A53T neurons. Conceptually, this stress-dominated state may be associated with high excitability that was measured at week 7 (Fig. 1e-g), while synaptic input lags (Fig. 1o-r).

In sharp contrast, the downregulated gene set indicated a broad energetic decline. KEGG terms showed depletion of “glycolysis/gluconeogenesis”, “fructose–mannose and pentose-phosphate pathways”, “HIF-1 and carbon metabolism”, and “amino-acid biosynthesis” (Fig. 3e). GO processes highlighted reduced “nucleotide phosphorylation”, “glycolytic flux”, “ATP generation from ADP”, and “pyruvate metabolism” (Fig. 3f), while GO components pinpointed downregulated “respiratory chain Complex I”, “inner mitochondrial membrane complexes”, and “mitochondrial protein–containing complexes” (Supp. Fig. 1d) -transcriptomic signature consistent with mitochondrial/oxidative phosphorylation (OXPHOS) insufficiency. GO molecular function analysis revealed significant downregulation of “amino acid transmembrane transporter activity”, “organic acid transmembrane transporter activity”, “DNA-binding transcription activator activity”, “collagen binding”, and “extracellular matrix structural constituent functions” (Supp. Fig. 1c). The downregulated network was centered with dysregulated hub genes related to metabolic-structural nexus, featuring nearly the entire glycolytic cascade (SLC2A1, PGK1, PKM, LDHA) and the primary cytoskeletal component ACTB as its most interconnected hubs (Fig. 3g). The volcano plot (Fig. 3a) mirrors this organization, with metabolic landmarks (e.g., HK2, PGK1, LDHA) shifted leftward (downregulated) and stress nodes (e.g., ATF4, DDIT3, SQSTM1, TFEB, NFE2L2) elevated -placing A53T mutant neurons in a metabolically compromised, low-ATP state.

Overall, our integrated electrophysiological and molecular findings define a unified timeline of neurodegeneration; as A53T mutant neurons move from early hyperexcitability (week 5; Fig. 1b-d) into mid-maturation (week 7), the induction of stress/apoptosis programs coincides with a downregulation of glycolysis and OXPHOS, a combination that may be associated with lower sEPSC frequency (Fig. 1o-q). This transcriptomic state provides the mechanistic insights into the mature neuronal functional inactivity (weeks 9–10) and may also underlie the observed response to EV treatment, which re-balances excitability and restores synaptic function (Fig. 4). A full list of DEGs and pathway analysis results is provided in Supplementary Table 1.

**Figure 4:**
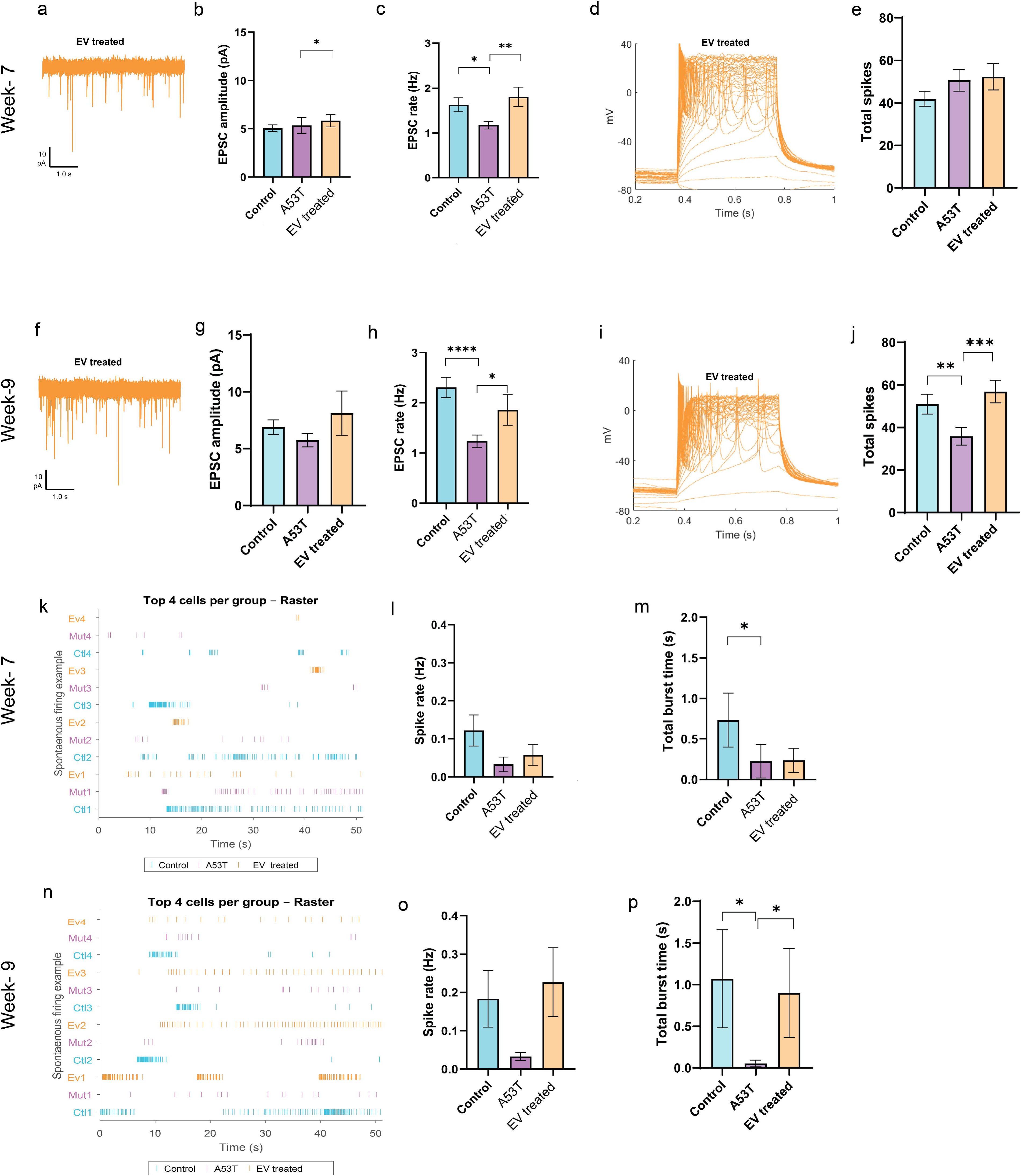
iPSC-derived EV Treatment Rescues A53T Neuronal Dysfunction. iPSC-derived EV treatment restores synaptic and intrinsic excitability deficits in A53T mutant neurons. **Week 7 rescue**: **(a)** Representative sEPSC traces from EV-treated A53T mutant neurons. EVs treatment significantly increases sEPSC **(b)** amplitude and **(c)** frequency compared to A53T mutant neurons. Representative **(d)** current-clamp traces and **(e)** total spike quantification with no significant differences across the conditions. **Week 9 rescue**: **(f)** Representative sEPSC traces from EV-treated A53T mutant neurons. Rescue effect of EVs treatment on sEPSC **(g)** amplitude showing a non-significant difference relative to untreated A53T mutant neurons and **(h)** frequency showing a significant effect. Representative **(i)** current-clamp traces and **(j)** total spike quantification showing complete functional restoration. Additional plots showing the spontaneous firing at two timepoints. **Week 7**: **(k)** Representative raster plots showing spontaneous action potentials in the four most active neurons per condition, accompanied by **(l)** average spike rate and **(m)** total burst time showing no rescue effect. **Week 9: (n)** Representative raster plots showing spontaneous action potentials in the four most active neurons per condition, accompanied by **(o)** average spike rate with no-significant rescue effect **(p)** total burst time with significant rescue effect. Data represent mean ± SEM with statistical analysis by the Mann-Whitney U test. (*P<0.05, **P<0.01, ***P<0.001).

### EVs treatment rescues electrophysiological and pathological abnormalities in A53T PD human models

Given the progressive functional decline observed in A53T mutant neurons and the demonstrated effectiveness of EVs in rescuing the deficits in both in vitro and in vivo ASD models from our previous work, we investigated whether healthy iPSC-derived EVs could ameliorate the emerging pathology. The EVs were administered to A53T mutant neurons and control lines starting at differentiation day 10 and applied every week throughout the differentiation period until the experiments were performed, as detailed in the Methods section. The treatment revealed promising early rescue effects by week 7, particularly in synaptic function, where deficits had become most apparent. At this timepoint, EV-treated A53T mutant neurons exhibited robust sEPSCs, with representative traces (Fig 4a) demonstrating restored synaptic activity comparable to untreated neurons (Fig 1p). EVs treatment produced a remarkable restoration of synaptic strength, with treated A53T mutant neurons exhibiting significantly increased sEPSC amplitudes compared to A53T mutant neurons (5.8 ± 0.6 pA in EVs-treated neurons vs. 5.3 ± 0.8 pA in A53T mutant neurons; P = 0.0485; Fig. 4b). This restoration was accompanied by enhanced synaptic connectivity, as evidenced by significantly increased sEPSC frequencies that were similar to control levels (1.8 ± 0.2 Hz vs. 1.6 ± 0.2 Hz in controls, P = 0.422), while untreated A53T mutant neurons had lower frequencies (1.2 ± 0.1 Hz), which was significantly increased by EV treatment (P = 0.0130; Fig. 4c). Representative evoked potential recordings from EV-treated A53T mutant neurons are shown in Fig. 4d; however, the number of evoked action potentials was not significantly different among the three groups (Fig. 4e).

From Weeks 9–10 (late maturation), when A53T mutant neurons display profound synaptic and firing deficits, most parameters of EVLtreated cultures showed functional activity that was similar to controls. Representative traces are shown in Fig 4f. No significant difference was observed with sEPSC amplitudes in treated compared to untreated A53T mutant neurons (Fig.4g). However, treated A53T mutant neurons maintained robust sEPSC frequencies (1.9 ± 0.3 Hz) that closely approached control levels (2.3 ± 0.2 Hz) (P = 0.7164) while remaining significantly elevated above A53T mutant neurons (1.2 ± 0.1 Hz; P = 0.0326; Fig. 4h). As shown with the representative evoked potential recordings from EV-treated A53T mutant neurons (Fig 4i), the restoration of spike generation was also remarkable, with EV-treated neurons producing significantly higher numbers of evoked action potentials compared to A53T mutant neurons (P = 0.0006; Fig. 4j).

To determine whether these improvements in intrinsic and synaptic properties translate into restored network function, we analyzed spontaneous firing and burst activity. At week 7, EV treatment did not result in a significant difference in spontaneous firing (Fig. 4k, l) and total burst time (Fig. 4m) between the treated and untreated groups. However, a composite electrophysiological maturation index spanning intrinsic, synaptic, and network metrics showed improvement of the EV-treated A53T mutant neurons toward the control trajectory without full normalization (Supp. Fig. 2a). Furthermore, a parameter heatmap highlighted the features most sensitive to the intervention at this stage -sEPSC frequency and spontaneous activity metrics (Supp. Fig. 2b). At week 9, spontaneous firing did not differ significantly across groups (Fig, 4n, o); however, total burst time was significantly reduced in A53T mutant neurons and increased following EV treatment to levels comparable to controls (P = 0.0225; Fig. 4p). During this timepoint, the composite maturation index and a parameter heatmap also displayed shared maturation profile between control and EV-treated neurons (Supp. Fig. 2a- c). Thus, the progressive enhancement from partial week 7 effects to complete week 9 rescue suggests that EV cargo requires time to integrate and exert maximal protective and trophic actions.

### Attenuation of pathological phosphorylated **α**-Syn accumulation in A53T DA neurons after EV treatment

Having shown that EV treatment robustly restores electrophysiological function in A53T DA neurons (Fig. 4), we next asked whether this rescue coincides with a reduction in pathogenic α-Syn burden. At the late maturation stage, we fixed and immunostained cultures for phosphorylated α-Syn (pSer129), the dendritic marker MAP2, and quantified pSer129 signal, detailed in the methods section. When normalized to the total intensity of DAPI, A53T mutant neurons showed a very significant accumulation of pSer129 compared to the control neurons (1.62 ± 0.25 in A53T mutant neurons vs. 0.12 ± 0.03; P =0.0003; Fig. 5a), and absolute pSer129 intensity was likewise markedly elevated in A53T mutant neurons 2.23 × 10L ± 0.39 × 10L AU (P < 0.0001; Fig. 5b). After treatment with EVs, both measures were dramatically reduced to 0.63 ± 0.11 (P = 0.0092 vs. A53T) and 1.2 × 10L ± 0.15 × 10L arbitrary units (AU) (P < 0.0001 vs. A53T), respectively. Finally, to account for changes in neuronal density and morphology, we also normalized pSer129 to the MAP2 signal, and MAP2 to DAPI. A53T mutant neurons exhibited significant elevation over control neurons (P = 0.0069; Fig. 5c), though MAP2 to DAPI normalization did not show a significant difference (Fig. 5d). Representative images are shown in Fig. 5e–g: controls display sparse pSer129 puncta amid dense MAP2-positive neurites, whereas A53T mutant neurons displayed marked accumulation of pSer129 aggregates, which appeared reduced following EV treatment.

**Figure 5:**
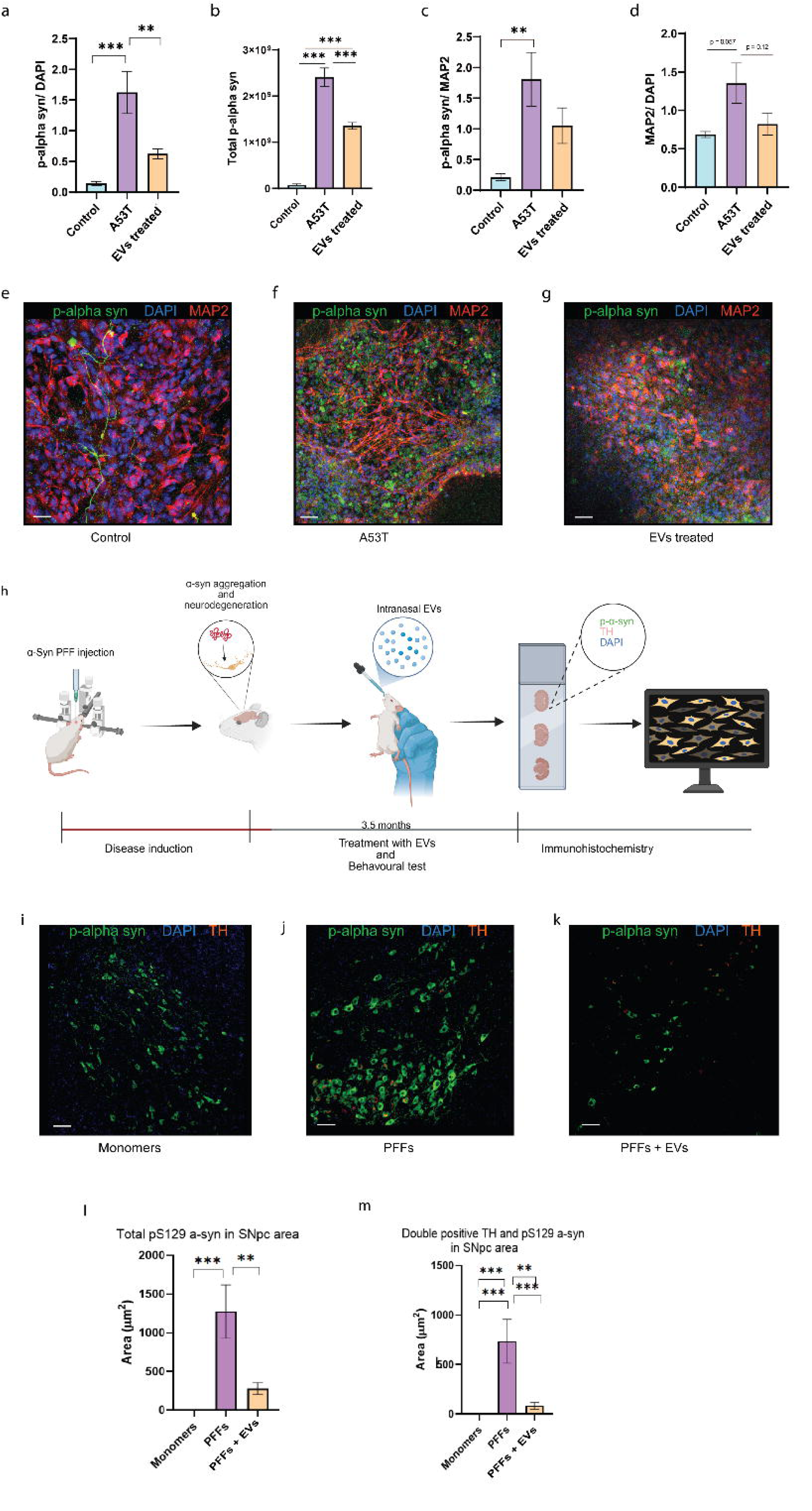
EVs Treatment Reduces Pathological α-Syn Accumulation. EVs treatment reduces pathological phosphorylated α-Syn accumulation in A53T DA neurons. Immunocytochemical analysis at day 60 using antibodies against phosphorylated α-synuclein (pSer129), MAP2 (dendritic marker), and DAPI (nuclei). **(a)** pSer129 normalized to DAPI intensity showing 13-fold increase in A53T mutant neurons with significant reduction following EV treatment. **(b)** Total pSer129 intensity measurements confirming its pathological accumulation in A53T mutant neurons and therapeutic reduction with EV treatment **(c)** pSer129 normalized to MAP2 signal accounting for neuronal density changes. **(d)** MAP2/DAPI ratio showing trends in dendritic coverage. Representative confocal microscopy images of **(e)** control, **(f)** A53T, and **(g)** EV-treated cultures. **(h)** Schematics showing the EVs treatment. Representative confocal microscopy images of a mouse brain slice under **(i)** monomer, **(j)** α-Syn PFFs, and **(k)** α-Syn PFFs with EV treatment. **(l)** Total pSer129 intensity confirming its pathological accumulation in the α-Syn PFF-injected group and therapeutic reduction after EV treatment **(m)** Quantification of TH/pS129 α-Syn double-positive area in the SNpc showing significant accumulation after α-Syn PFF treatment and reduction following EV treatment. pSer129 (green), MAP2 (red), and DAPI (blue). Scale bars are 50um. Data represent mean ± SEM. One-way ANOVA with Tukey’s multiple comparisons test (*P < 0.05, **P < 0.01, ***P < 0.001).

To examine whether these effects extend *in vivo*, we analyzed substantia nigra pars compacta (SNpc) sections from α-Syn PFF-injected mice with or without intranasal EV treatment, as displayed in Fig. 5h. Quantification revealed a marked accumulation of pS129 α-syn in the SNpc of α-Syn PFF-injected animals compared to monomer controls (Fig. 5i–k show representative examples). EV treatment significantly reduced total pS129 α-syn area (P = 0.0002; Fig. 5l) as well as the area of double-positive TH/pS129 α-syn neurons (P = 0.0006; Fig. 5m) compared to those only with α-Syn PFF-injection, indicating suppression of α-syn aggregation *in vivo*. Together, these results show that EVs not only restore intrinsic and synaptic function *in vitro* but also mitigate α-syn pathology in both cellular and animal models of PD, providing a mechanistically coherent rescue of A53T DA neuron health. To further address the translational relevance, we conducted a pilot behavioral assessment on α-Syn PFF-injected mice that were treated with EVs intranasally (Supp. Fig. 3). As shown in Supp Fig. 3a, the experiment consisted of α-Syn PFF injection, EV treatment, and subsequent motor testing. At 3.5 months post- α-Syn PFF injection, body weight was comparable among the groups (Supp. Fig. 3b). In the motor coordination assay performed 1.5 months post- α-Syn PFF injection, latency to fall did not differ significantly between the α-Syn PFF-injected group and EV-treated group (Supp. Fig. 3c). However, at 3.5 months post- α-Syn PFF injection, EV treatment significantly improved motor performance relative to the α-Syn PFF-injected group (Supp. Fig. 3d). Overall, despite the limited sample size, our preliminary data showed trends toward preserved motor performance following EV administration, consistent with the *in vitro* rescue profile.

## Discussion

Our study extends previous findings of altered synaptic activity in PD patient-derived DA neurons during weeks 7–9 of differentiation (40–42, 50–52). However, whether these changes evolve linearly with maturation and how intrinsic excitability and network activity change over time, has remained unclear in our previous research. Here, we uncover a biphasic course of A53T SNCA mutant neurons, showing early hyperexcitability, followed by synaptic failure and late hypoexcitability, that mirrors emerging concepts of prodromal circuit disruption in synucleinopathies. We then map week-7 as a molecular inflection point characterized by activation of stress/apoptotic programs along with coordinated suppression of glycolysis and OXPHOS. We further show that healthy iPSC-derived EVs can reverse both functional and molecular pathology of A53T mutant neurons, as evidenced by restoration of intrinsic excitability, synaptic transmission, spontaneous network activity, and reduced phosphorylated α-Syn accumulation.

The biphasic electrophysiological phenotype identified in A53T DA neurons, characterized by an early phase of hyperexcitability followed by progressive transition to hypoexcitability and synaptic failure, adds temporal insight to synucleinopathy progression and constitutes one of the key findings of our study. This pattern aligns with emerging evidence that network over-activity represents an early and potentially pathogenic event in PD, preceding overt neuronal dysfunction and death (53, 54). For instance, patient-specific LRRK2 neuronal networks have been reported to show elevated bursting without overt neurodegeneration, supporting the concept that early hyperactivity is pathogenic rather than epiphenomenal (53). Early network hyperexcitability has also been observed in α-Syn transgenic mice, where hippocampal circuits display aberrant bursting before cognitive impairment, further strengthening the idea that hyperactivity is an upstream driver in synucleinopathies (55, 56). This hyperexcitable state plausibly imposes metabolic load and Ca²L stress, consistent with excitotoxic frameworks and DA-neuron selective vulnerability models that link sustained activity and Ca²L entry to bioenergetic strain and oxidative stress in SNpc neurons (57–59). By contrast, during the late phase, we observed reduced firing and diminished sEPSC frequency, mirroring the connectivity loss seen in PD patient iPSC models across genotypes (including SNCA, PRKN, LRRK2, and sPD), in which synaptic event rates are consistently altered at 7-9-week differentiation stages (40, 41). In general, our biphasic trajectory resonates with other neurodegenerative disorders such as Alzheimer’s disease, where network hyperexcitability is now recognized as an early hallmark that can precede and predict later synaptic loss and cognitive decline (60–62). This comprehensive temporal analysis reveals that A53T α-Syn imposes an evolving pattern of synaptic dysfunction that progresses from selective connectivity deficits to comprehensive network failure. The early preservation of synaptic strength despite reduced connectivity suggests initial compensatory mechanisms that ultimately prove insufficient to prevent the severe deterioration observed at later stages.

To understand what drives the transition from early over-activity to late failure, we profiled transcriptomes at week 7, when electrophysiological divergence becomes prominent. We found a broad transcriptional reorganization with 1,190 DEGs, suggesting a molecular basis for the electrophysiological changes. Among upregulated genes, enrichment analyses revealed a coordinated activation of cell-stress and pro-death pathways, including p53, JAK–STAT, FoxO, TNF signaling, as well as apoptosis and cytokine–receptor interactions. This pattern suggests that neurons at this stage are engaged in an integrated stress response, shifting toward inflammatory and pro-apoptotic signaling as functional decline emerges. GO terms reinforced this signature, highlighting activation of endoplasmic reticulum (ER) stress-induced apoptotic signaling, autophagy regulation, and inflammatory response pathways. These findings point to an integrated stress response involving nuclear chromatin remodeling and cytosolic stress granule formation, with concurrent extracellular matrix remodeling, suggesting propagation of stress effects to the tissue level. Overall, this transcriptional state is consistent with a model in which sustained hyperexcitability, observed from our electrophysiological findings, is associated with a transition toward proteostatic stress, inflammation, and cell death pathways.

In contrast, the downregulated gene set revealed a pronounced metabolic deficit. KEGG analysis showed broad suppression of glycolysis/gluconeogenesis, the pentose phosphate pathway, HIF-1 signaling, amino acid biosynthesis, and central carbon metabolism. This was supported by GO biological process enrichment, which indicated reduced glycolytic flux, ATP generation, and pyruvate metabolism. Consistently, GO cellular components analysis pointed to impairment of the mitochondrial electron-transport machinery, including oxidoreductase complexes and respiratory chain Complex I, indicating disruption of oxidative phosphorylation. Molecular function analysis further showed a decline in amino acid and organic acid transport across membranes, pointing to a broader disruption in cellular metabolic balance. Together, these data align with reports that α-syn proteopathy destabilizes mitochondria, reduces ATP production, and promotes fragmentation and mitophagy in DA neurons (58), as well as iPSC studies linking α-syn accumulation to bioenergetic dysfunction and ER stress (63).

Mechanistically, the electrophysiological changes can be interpreted in the context of this molecular architecture: early hyperactivity elevates energetic demand, but by week 7, the neurons activate stress and apoptotic programs as glycolytic and oxidative phosphorylation capacity declines. This energy demand-supply mismatch is accompanied by reduced presynaptic release probability and impaired vesicle recycling, reflected in decreased sEPSC frequency and intrinsic excitability. The concomitant downregulation of cytoskeletal modules (e.g., ACTB hub) may further destabilize synaptic architecture and impair bursting. In summary, week 7 marks a tipping point at which stress signalling rises while metabolic capacity declines.

Delivering healthy iPSC-derived EVs produced a time-dependent rescue, partial by week 7, and complete by week 9. The rescue involved intrinsic, synaptic, and network domains, accompanied by reduced pSer129 accumulation. EVs carry proteins, lipids, and regulatory RNAs that can modify the state of recipient cells (31). In line with this, in our previous study, we demonstrated that iPSC-derived EVs are enriched in plasticity and homeostatic regulatory cargoes (36). These cargoes may underlie their ability to drive a time-dependent rescue of functional abnormalities in A53T mutant neurons. Engineered EVs have been used to deliver antisense oligonucleotides or siRNA against SNCA, leading to reduced total and pSer129 and amelioration of pathological features in PD mouse models (64, 65). Furthermore, intranasally administered stem-cell EVs were shown to exhibit high brain infiltration and therapeutic effects across neurological models (66). In the present study, the EVs delivered via this route successfully rescued the abnormal behavioural phenotypes in α-Syn PFF-injected PD models. Notably, we have also previously applied the same EV preparations in ASD mouse models, where they similarly rescued disease-related phenotypes (36). Collectively, these findings indicate that iPSC-derived EVs have the potential to attenuate PD-related phenotypes in both *in vitro* and *in vivo* models.

While the active cargo remains undefined in our study, the delayed recovery kinetics, progressing from partial to full over ∼2 weeks, suggest a programmatic reset rather than a transient modulatory effect on ion channels. A plausible model is that iPSC-derived EVs attenuate stress and apoptosis signalling pathways, while reactivating metabolic functions and enhancing proteostasis. Together, these changes may break the feed-forward loop by uncoupling neural stress responses and α-syn aggregation and propagation. This framework is consistent with prior iPSC-based PD studies demonstrating synaptic deficits across genotypes and convergent transcriptomic alterations, reinforcing the need for multi-target therapeutic strategies rather than single-node intervention (40, 67). Overall, the robust and sustained reversal of A53T pathology by iPSC-derived EVs underscores their therapeutic promise for synucleinopathies, acting to counteract both the stress-inflammation activation and metabolic shutdown elucidated in our transcriptomic analyses.

In conclusion, our data support a mechanistic cascade in A53T DA neurons in which early hyperexcitability elevates energetic demand, and by week 7, neurons enter a stress and apoptosis program as glycolysis and OXPHOS falter, precipitating synaptic failure and intrinsic regression. Timed delivery of healthy iPSC-derived EVs interrupts this cascade, rebalancing excitability, restoring synaptic transmission, and reducing pSer129 aggregation. In light of convergent evidence across PD and AD that network hyperexcitability is an actionable early node, our findings argue for network-centric, multimodal interventions with iPSC-derived EVs offering a clinically plausible, cell-free platform that can be native or engineered to reprogram stressed neurons and stabilize vulnerable circuits. Since our current findings are based on A53T mutant neurons, future work should further validate the phenotype and EV-mediated rescue across multiple PD genotypes and sporadic PD lines to establish wider applicability. More mechanistic studies, such as targeted proteomic/miRNA profiling, loss-of-function approaches, orthogonal metabolic assays, *in vivo* validation in larger cohorts, and post-EV transcriptomic analyses, will be required to pinpoint the major EV effectors and link molecular alterations to functional recovery.

## Supporting information

Supplemental Figure 1: Dysregulated Metabolic Pathways in A53T mutant neurons

Supplemental Figure 2: Composite maturation index and electrophysiological parameter heatmaps

Supplemental Figure 3: In Vivo Behavioral Rescue by EV Treatment

---

**Supplementary Figure 1: Dysregulated Metabolic Pathways in A53T mutant neurons**

Gene Ontology and KEGG pathway analysis of 1190 DEGs from RNA-seq analysis (week 7). **(a)** GO Molecular Functions demonstrating increased DNA-binding TF activity, cytokine-receptor binding, protein-kinase binding, and more dysregulated pathways. **(b)** GO Cellular Components showing upregulation of nuclear chromatin, mitochondrial matrix, and cytosolic stress granules, consistent with transcriptional re-programming, organellar strain, and proteostasis pressure. **(c)** GO Molecular Functions demonstrating loss of carbohydrate kinase activity and oxidoreductase functions. Analysis reveals systematic collapse of energy production and structural maintenance systems in A53T mutant neurons**. (d)** GO Cellular Components showing downregulation of respiratory chain complexes and mitochondrial machinery. Analysis based on n=3 control and n=3 A53T cell lines with 2 biological replicates each. DEGs were defined as those with an FDR < 0.05 and (**|**log_2_ fold change |) > 0.58 (≈1.5-fold).

**Supplementary Fig. 2. Composite maturation index and electrophysiological parameter heatmaps**

**(a)** Electrophysiological maturation index integrating intrinsic, synaptic, and network properties at weeks 7 and 9. A53T mutant neurons show divergence from control trajectories, while EV-treated neurons realign toward the control profile by week 9. **(b)** Heatmap of individual electrophysiological features at week 7 highlights partial rescue in synaptic measures but incomplete correction of spike threshold. **(c)** The heatmap demonstrates broad normalization across intrinsic, synaptic, and network metrics in EV-treated neurons at week 9.

**Supplementary Figure 3: *In Vivo* Behavioral Rescue by EV Treatment**

EV treatment ameliorates motor deficits induced by α-Syn PFF *in vivo*. **(a)** Adult C57BL/6 mice received intrastriatal injections of α-Syn monomers (control), α-Syn PFFs, or α-Syn PFFs plus intranasal EV treatment (4 μL containing 2-3×10¹L particles/μL every other day for 3.5 months). **(b)** Body weight monitoring showed no systemic toxicity across treatment groups. **(c)** Inverted screen test at 1.5 months post-injection showing trends toward motor impairment in α-Syn PFF-treated mice. **(d)** Inverted screen test at 3.5 months demonstrating significant rescue of motor deficit by EV treatment. Data represent mean ± SEM. Final group sizes: monomers n=2, α-Syn PFFs n=2, α-Syn PFFs+EV n=4 (reduced due to planned tissue collections). Statistical analysis by appropriate tests for small sample comparisons.

