## Supplementary figures and images for "iPSC-derived extracellular vesicles rescue deficits in human and mouse models of Parkinson’s disease"

### Supplemental Figure 1: Dysregulated Metabolic Pathways in A53T mutant neurons

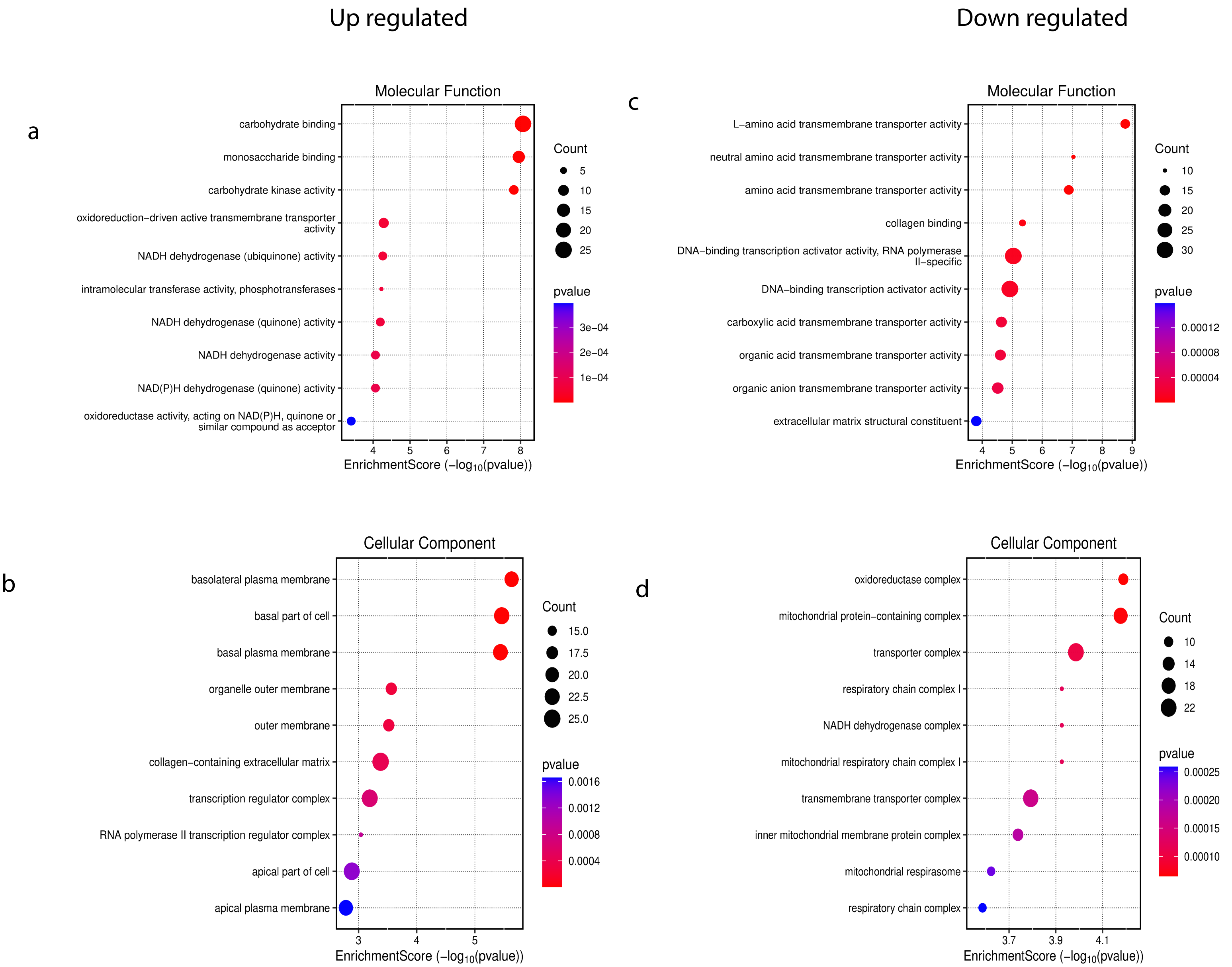

### Supplemental Figure 2: Composite maturation index and electrophysiological parameter heatmaps

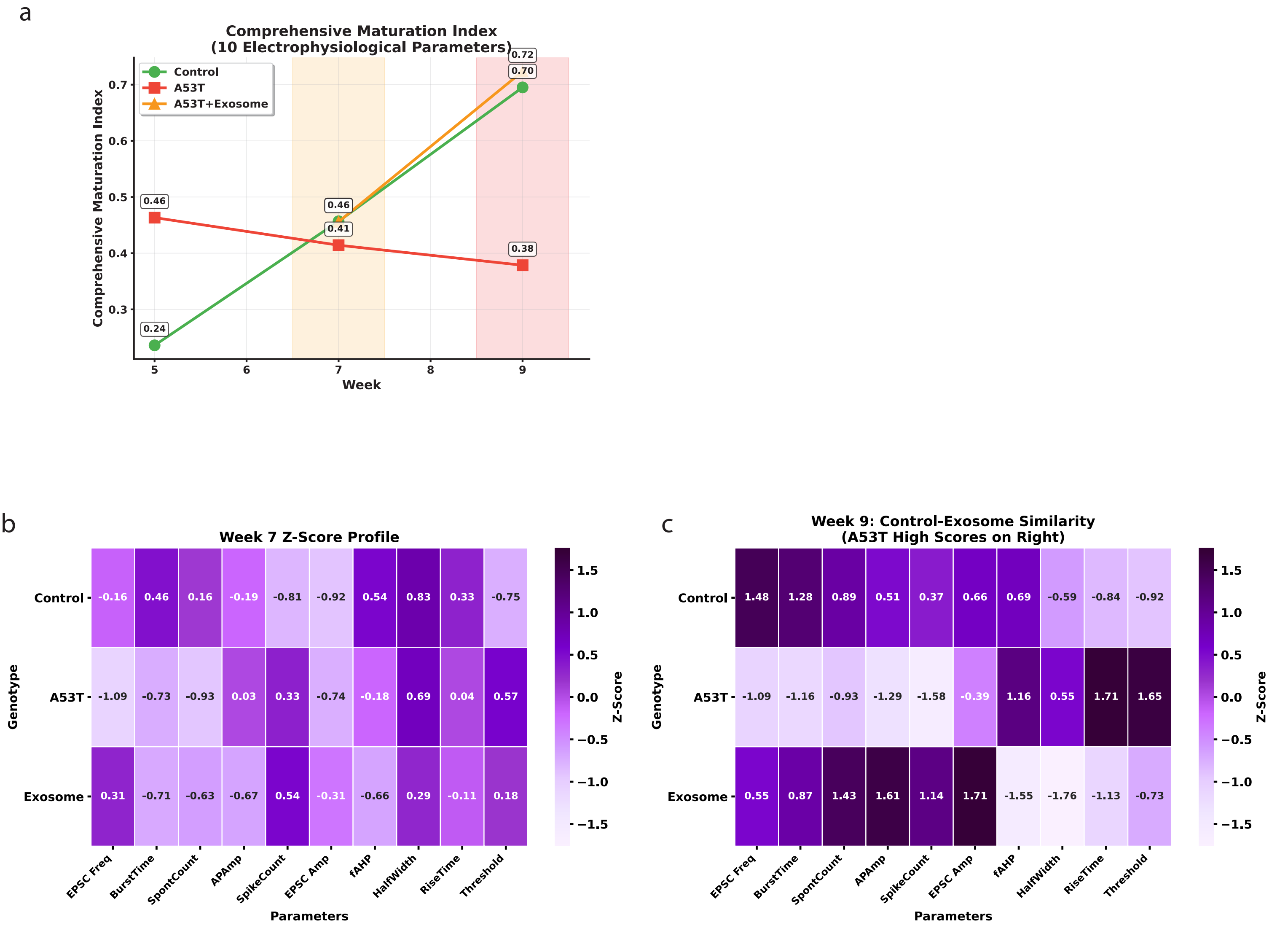

### Supplemental Figure 3: In Vivo Behavioral Rescue by EV Treatment

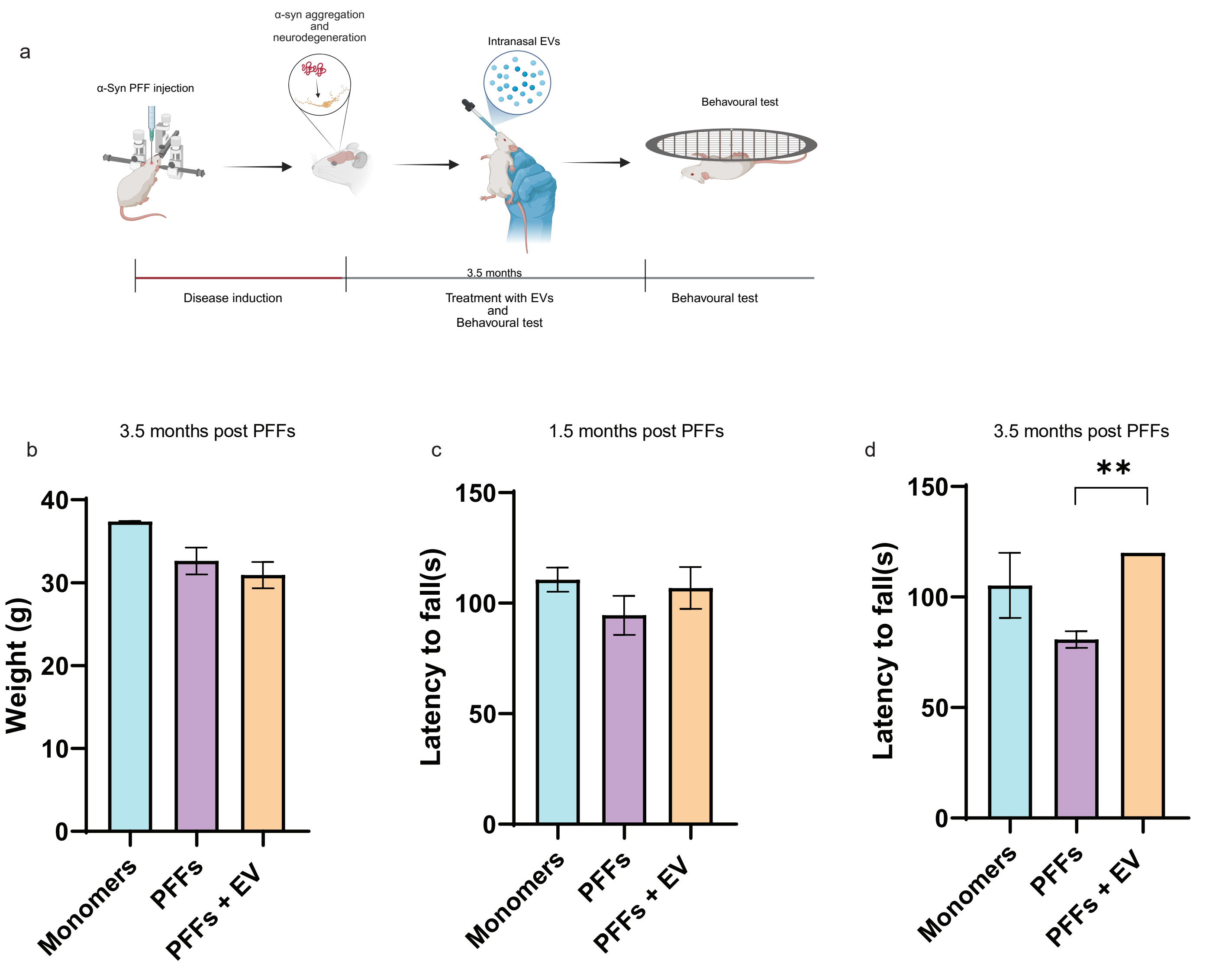
